# Nutrition mediates extreme growth variation through deep changes in gene expression in the water strider *Microvelia longipes*

**DOI:** 10.64898/2026.08.27.747451

**Authors:** Ingrid Dourlens, Séverine Viala, Kiran Padmanabhan, Abderrahman Khila

**Affiliations:** Institut de Génomique Fonctionnelle de Lyon, CNRS UMR 5242, Ecole Normale Supérieure de Lyon, Université Claude Bernard Lyon 1, 46 allée d’Italie, 69364 Lyon Cedex 07, France

**Keywords:** phenotypic variation, comparative transcriptomics, growth exaggeration, histone modification, gene function

## Abstract

Exaggerated sexually selected traits are known to be highly variable and their degree of expression is dependent on nutritional input. Yet the molecular mechanisms linking nutritional variation to phenotypic variation remain poorly understood. Here, we investigate how nutritional input shapes the development of male rear leg length, an exaggerated and highly variable trait in the water strider *Microvelia longipes*, using comparative transcriptomics and RNA interference gene knockdown experiments. We demonstrate that nutrition is the primary driver of gene expression variation, with male exaggerated rear legs exhibiting the highest number of nutrition-responsive genes. Moreover, the increase in morphological divergence between leg types or sex, which is systematically exacerbated by rich nutrition, is associated with increased number of leg-biased genes. These comparative analyses allowed us to identify BMP11 as specifically enriched in female and male rear legs. Knockdown of BMP11 abolishes nutritional plasticity in leg length only in males, positioning it as a key integrator of environmental, sex and developmental signals. Our findings reveal that transcriptional modulation provides a molecular interface between nutrition and trait exaggeration. This work advances our understanding of how environmental cues are translated into complex phenotypes and highlights the role of developmental plasticity as a substrate for evolutionary change.

## Introduction

Phenotypic variation is widespread in nature and represents a major pillar of evolutionary change [1]. Yet, our understanding of the molecular mechanisms responsible for generating phenotypic variation during development remains limited [2–5]. Exaggerated secondary sexual traits represent some of the most astounding examples of phenotypic variation [6].

These traits are also known to be plastic, and their degree of expression is heavily influenced by nutritional input [7–11]. As such, exaggerated secondary sexual traits represent powerful models to investigate how the interaction between genetic, epigenetic, and environmental factors generates high phenotypic variation among individuals of the same population.

Variation in gene expression provides an important molecular interface through which environmental conditions can influence the phenotype [12–14]. This is particularly relevant for exaggerated sexually selected traits, whose growth is often considerably more sensitive to nutrition than that of other tissues within the same individual. Studies in beetles have shown that this increased condition-dependence involves the nutrition-dependent regulation of developmental and growth pathways, including insulin/insulin-like growth factor signalling (IIS), as well as sex- and tissue-specific regulators [15–17]. For example, in the dung beetle *Onthophagus taurus*, the sex-determination gene *doublesex* contributes to the sex-specific development of exaggerated horns, while the growth inhibitor FoxO contributes to suppressing the growth of the horn under poor nutritional conditions [16, 18]. These findings illustrate how nutritional sensitivity of exaggerated traits can emerge through differential regulation of growth programs across sexes, tissues and nutritional environments.

Most of our understanding of these mechanisms, however, comes from holometabolous insects, particularly beetles, exhibiting mostly discrete or nonlinear weapon development. In several of these systems, nutrition-dependent growth produces alternative male morphs through threshold-like developmental responses [9, 19]. Yet many exaggerated sexually selected traits vary continuously with body size and nutritional condition. Whether such continuously variable traits rely on similar molecular mechanisms, and more generally how the degree of transcriptional variation relates to the degree of morphological exaggeration, remains poorly understood. Expanding these analyses to continuously variable traits and to hemimetabolous insects therefore provides an opportunity to determine whether common principles underlie the evolution and developmental plasticity of exaggerated structures across divergent developmental systems.

Here we address these questions in the water strider *Microvelia longipes* [20, 21]. This species exhibits striking variation in the size of the rear legs both within and between the sexes [21, 22]. The legs are significantly longer in males compared to females, but also exhibit extreme variation among males of the same population. The length of the rear legs in males is hyperallometric and scales disproportionately to body size with a slope coefficient exceeding 3; one of the highest known [21, 22]. Males use their rear legs as weapons during contests, and long-legged males have a significantly higher chance to win the contest, dominate egg-laying sites, and sire the majority of eggs laid by the females that visit the sites [21]. Male rear leg size variation is highly responsive to nutrition as reaction norm experiments, using isogenic lines with depleted genetic variation through brother-sister inbreeding, have shown that individuals with the same genetic background will grow long or short rear legs in rich and poor nutritional conditions, respectively [21, 23]. While we know that this variation is heavily influenced by nutrition, the mechanisms mediating this effect of nutrition on leg growth remain unknown.

To better understand the interaction between molecular and phenotypic variation, we conducted an unbiased genome-wide examination of changes in gene expression induced by nutritional variation. Using an inbred line, we examined the effect of nutrition on nymphal development in both males and females. We produced the transcriptomes of the developing legs in these nutritional treatments to uncover how nutrition alters gene expression for the first time in this species. We identified and functionally tested genes whose expression is nutrition-dependent in both sexes and that show a role in regulating nutrition-dependent leg growth variation specifically in males.

## Material & methods

### Animal husbandry and nutritional treatment

Inbred lines were generated from a natural population collected in French Guyana in March 2023 as described in previous study [21]. We used long-legged isogenic line for further experiments, as its phenotypic variation is the most marked. The bugs were maintained in the laboratory at 27-29°C and 50% humidity in water tanks and fed on crickets. For the nutritional experiment, first instar nymphs were collected just after hatching in a 7 hours windows, and 50 individuals were fed with 1 fresh cricket until they reach in the second nymphal instar to avoid mortality. Then we fed nymphs in either poor or rich nutritional condition. Individuals from the same nutritional condition were raised in the same water tank. In the poor condition, we used only 1 frozen rear leg to feed 20 second instar or 50 third to fifth nymphal instar bugs. In rich condition, 20 second instar nymphs or 10 older nymphs were fed with 1 fresh cricket. We hypothesized that molecules involved in extreme growth variation in response to nutrition are actively expressed during the final nymphal instar, a critical stage preceding the burst of growth. Then we selected the males and females in the final nymphal instar biased on their absolute rear leg length, since the differences between nutritional diet were already visible in male last nymphal instar (**Supplementary Figure S1**). Selected individuals were enriched in short-legged or long-legged individuals either in poor or rich nutrition, and were used for RNA extraction (**Supplementary Figure S2**).

### Statistical analyses and leg measurements

All statistical analyses were performed in RStudio 4.3.0. The number of individuals used for measurement per condition is in Supplementary Table 1. We measured the tibia from the rear legs that is proportional to the absolute rear leg length, with a VHX 7000 microscope (Keyence) and the VHX 7000 software (version 1.3.11.2).

The effects of RNAi treatment, nutrition, and their interaction on rear-leg length and body size were tested using two-way ANOVAs. Model assumptions were assessed by visual inspection of residual diagnostic plots. Moreover, complementary non-parametric analyses yielded qualitatively similar conclusions, supporting the robustness of the results obtained with the parametric models. Body size-corrected rear-leg length was obtained from the residuals of a linear regression of rear-leg length against body size and analyzed using the same two-way ANOVA framework. Standardized effect sizes and 95% confidence intervals were calculated to compare treatment and nutritional effects across conditions.

### Tissue harvesting and RNA sequencing

We collected the rear legs from both sexes and the forelegs from male 5^th^ nymphal instar nymphs (2 days after molting) that were raised under to the two nutritional conditions described above. The three replicates of each condition (nutrition, sexes and legs) correspond to a pool of individuals (**Supplementary Table S1**). The dissection of the pairs of legs, dissociated from the thorax, was performed in RNAse free 0,1% PBS-Tween using fine needles, each pair of legs was incubated immediately on ice in tubes and was kept at -70°C until we collected all the samples. Legs were grinded in liquid nitrogen and immediately tubes were filled with TRIzol (Invitrogen). RNA extractions were performed according to manufacturer protocol. The concentrations were assessed using the Qubit 2.0 Fluorometer (Invitrogen). Library construction and sequencing were performed by SNP&SEQ Technology Platform in Uppsala. The samples were sequenced using Illumina NovaSeq X Plus sequencing technology with a paired-end read length of 150 bp.

### Mapping

Technical replicates were produced to increase read depth, and were analyzed separately in this part. Raw RNA-seq reads were trimmed with Trimmomatic v. 0.39 [24]. Specifically, reads were trimmed if the sliding window average Phred score over four bases was < 15, and only reads with a minimum length of 36 bp. Read quality was assessed before and after trimming using FastQC (v. 0.11.9, https://www.bioinformatics.babraham.ac.uk/projects/fastqc/) and aggregated using MultiQC v. 1.12, [25]. Braker annotation was used as a reference for read alignment and transcriptome quantification. We removed around between 9 and 55% of ribosomal RNA using SortMeRNA v. 4.3.6, [26] with rRNA sequences from different Gerris species as reference. We used the filtered reads to map against *Microvelia longipes* genome and we obtained around 77% of uniquely mapped reads using STAR method (v. 2.7.8a (Dobin et al., 2013) with default settings. Only reads with good alignments were kept for further analyses using samtools v. 1.21, [27] to filter out reads with a map quality below 255.

This last output was used for the creation of count tables associated with a number of read per “gene” using featureCounts v. 2.0.1, -p -C -B settings [28]. Concatenated bam files from technical replicates were used to make the estimation of transcript abundance in FPKM with StringTie v2.1.4, [29]. Details about quality and mapping score are found in **Supplementary Table S1**.

### Identification of nutrition-, sex- and leg-biased genes

The transcriptomic approach was performed with three levels of comparisons, nutrition, sex and the leg type (**Supplementary Figure S3**). The comparative analysis was performed as described in [30]. In brief, genes were retained if their mean FPKM exceeded 2 in at in at least one tissue- and sex-specific group under either poor or rich nutritional conditions. Raw counts from the retained genes were subsequently used for differential expression analyses with DESeq2 [31]. DEseq2 script will be soon available from Dryad Digital Repository.

Three differential expression analyses were performed between homologous tissues from individuals raised under poor or rich nutrition, in order to find nutrition-biased genes in male rear legs, male forelegs or female rear legs. Two differential expression analyses were performed between male and female rear legs to find sex-biased genes in poor or rich nutrition. Finally, two differential expression analyses were performed between male fore and rear legs to find leg-biased genes in poor or rich males. Technical sequencing replicates were first collapsed using the collapseReplicates() function, which sums raw counts across sequencing runs for each biological sample. The DEseq2 model accounted for sequencing platform and biological replicate effects. Genes were considered differentially expressed when they showed an adjusted p-value (Padj) < 0.05 and an absolute log2 fold change > 0.58, corresponding to a fold change > 1.5.

### Interaction between nutrition and sex or leg gene expression

In order to detect a possible interaction between nutrition and sex or leg regulation, we combined our list of nutrition-biased genes with the list of either sex- or leg-biased genes. Using Fisher’s exact tests, we identified enrichment of genes with both nutrition-biased expression and either sex- or leg-biased expression (**Supplementary Table S2**).

### Gene Ontology analysis

Gene names and functions were annotated by sequence similarity against the NCBI “nonredundant” protein database using Blast2GO [30]. The Blast2GO annotation was then provided to detect Gene Ontology terms enrichment (p-value < 0.05) using the default method of TopGO Rpackage version 4.3.0.

### Nymphal RNAi and nutritional treatment

Double-stranded RNA (dsRNA) was produced and purified for *BMP11* and *Ubx* as described in [22]. The purified dsRNA was eluted in Spradling injection buffer (Rubin and Spradling, 1982) at a concentration of 5 µg/µL for *BMP11* and 0.5 or 5 µg/µL for *Ubx*. Nymphal injections were performed in the isogenic line selected for long-legged males [21] at the third instar as described in [32]. We injected nymphs with the dsRNA of the targeted gene corresponding and, in parallel, with buffer as negative control in two different experiments for *Ubx* and *BMP11* RNAi. Directly after injection, *Ubx*, *BMP11* and negative control nymphs were fed with standard nutrition for overnight to recover. Nymphs injected with dsRNA or Buffer were treated in parallel with the nutritional conditions described above, the day after the injection. We injected nymphs with two non-overlapping fragments of the BMP11 gene and we did not see differences in the ratio between male rear leg length and body size (**Supplementary Figure S4**). For other candidate genes tested, see **Supplementary Table S3**.

## Results

### Nutrition modulates rear leg growth in the last nymphal instar

To determine the link between gene expression and extreme growth variation in response to nutrition, we raised individuals under poor or rich diet and analyzed gene expression profiles using comparative transcriptomics. *M. longipes* male rear legs become excessively long after a burst of growth that occurs between the last nymphal instar (i.e. the 5^th^ instar) and adulthood, such that we can already discriminate small-legged from long-legged males at the 5^th^ nymphal instar (**Supplementary Figure S1**).

We found that fifth instar males fed on rich diet had significantly longer rear legs compared to starved males, with an increase of about 22% in mean leg length (1138µm and 934,9µm in rich and poor condition respectively, W = 25876, p< 2.2e-16, **Figure 1A**). Leg length was only 7,9% higher in females raised on rich compared to females raised on poor food (W = 34932, p-value < 2.2e-16, **Figure 1B**), consistent with the low phenotypic variation in females. This result shows that nutrition influences male rear leg growth variation in late-nymphal instar. These data reinforce the hypothesis that the genes involved in extreme growth variation are expressed in the rear legs of the fifth instar nymphs, and that variation in the levels and profiles of expression of these genes will modulate the observed variation in leg length.

**Figure 1.**
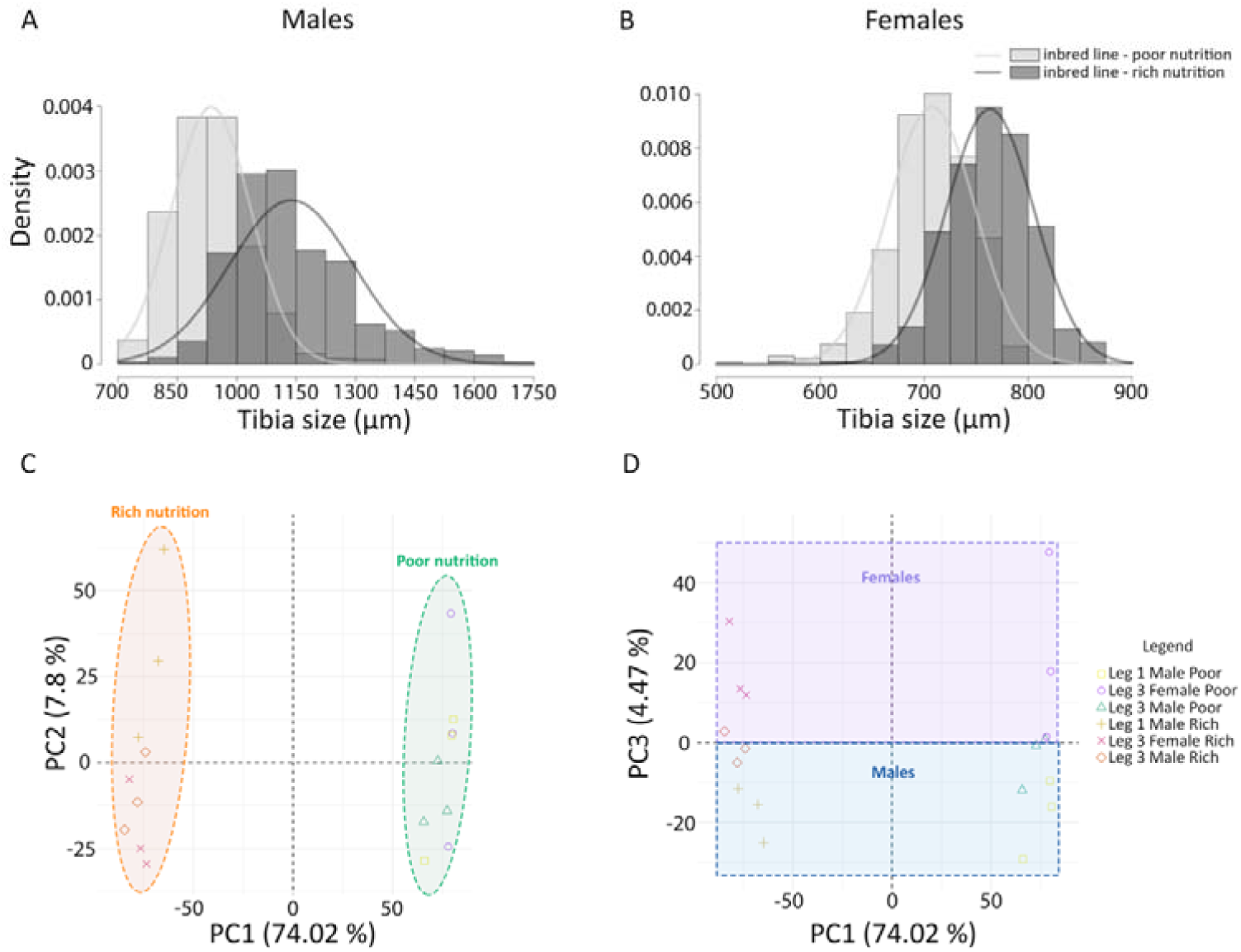
**A,B** Variation in rear tibia length in (A) males and (B) females in the last nymphal instar. The nymphs have been raised under rich (dark grey) or poor nutrition (light grey). **C, D** Principal component analysis (PCA) on the whole transcriptomic data of male forelegs and rear legs and female rear legs raised under poor or rich nutrition. **C** First axis (PC1) explains differences between nutritional treatments while PC2 explains differences between leg type (**Supplementary Figure S5**). **D** PC3 explains differences between sexes.

To test how nutritional variation contributes to extreme growth variation, we compared transcriptional profiles of rear legs from short-legged and long-legged males raised on poor or rich nutrition, respectively. Male forelegs and female rear legs under both nutritional treatments were added as comparisons in this analysis because they covary in an isoallometric pattern with body [21, 22, 30]. Of note, the main difference between these three leg tissues is length, at the exception of male forelegs which carry a sex comb [33]. A Principal component analysis (PCA) on these datasets revealed that nutritional treatment accounted for 77,3% of the variation in gene expression as the first principal component (PC1, x-axis) separated the sample based on diet (**Figure 1C**). The second major axis (PC2) separated the samples by tissue type and explained only 6,86% of the variation between various leg tissues (**Supplementary Figure S5**). The third major axis (4,17% of the variation) separated the sexes (**Figure 1D**). Together, these results show that nutrition represents the primary source of variation in gene expression, followed by serially homologous tissues then sex.

### Tissue experiencing the most exaggerated growth is enriched in nutrition-biased and leg-biased genes

The rear legs of males undergo the most striking exaggeration of growth on rich relative to poor treatment, and this growth variation could be associated with changes in the transcriptional landscape (**Figure 1C**). We wanted to test whether rear legs of males raised on rich diet respond more intensely to nutrition than other legs in males or females. We therefore compared transcriptional profiles of nutrition-dependent genes across the legs of males and females (i.e. genes whose levels of expression change in homologous legs with diet). Our data show that almost 7 times more genes responded to differences in nutrition compared to differences in tissue or sex, consistently with the high proportion of variation explained by nutrition in PCA (**Figure 1C**, **Table 1**). Rich-biased genes (genes significantly upregulated in rich diet) are more abundant than poor-biased genes regardless of leg type (**Figure 2A**, **Table 1**). Furthermore, the overall magnitude of differential gene expression, illustrated by log2 fold change, is more intense under rich nutrition in each tissue (**Figure 2C**). Together, these results suggest that either rich nutrition induces a strong upregulation of several genes in legs of rich-fed individuals, and/or poor nutrition can lead to a strong downregulation in legs of poor-fed individuals in *Microvelia longipes*.

**Figure 2.**
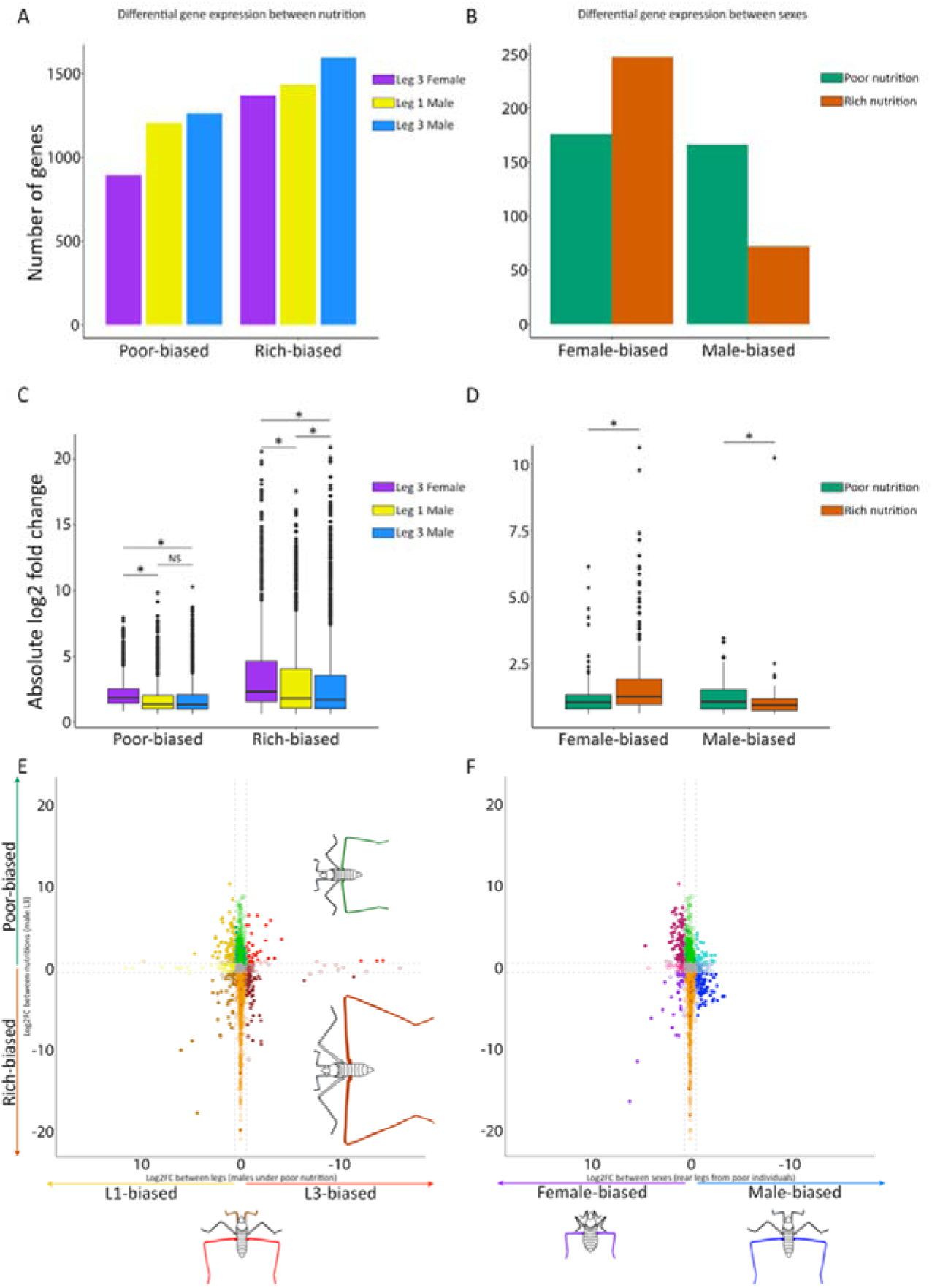
**A-D** Gene expression magnitude is represented through (A, B) differences in absolute log2 fold change (Wilcoxon) and (C, D) the total number of genes that are (A, C) nutrition- or (B, D) sex-biased. **A, C** Nutrition-biased genes are either up-regulated in poor or rich nutrition (poor-biased and rich-biased respectively) in each of the respective tissues: female third legs, male forelegs and male rear legs (purple, yellow and blue respectively). **B, D** Sex-biased genes are either up-regulated in females’ or males’ rear legs (female-biased and male-biased respectively) from individuals raised under poor or rich diet (green and orange respectively). Significant biased-genes display abs(log2 fold change)>0.58 and adjusted p-value <0.05. **E, F** Comparison between nutrition-biased genes in male rear legs and (E) leg-biased genes in poor males) or (F) sex-biased genes in poor individuals. Y axis represents the same genes in the two panels: poor- and rich-biased genes that are illustrated in green and orange respectively. X axis indicates either (E) L1- and L3-biased genes that are illustrated in yellow and red respectively or (F) female- and male-biased genes that are illustrated in purple and blue respectively. Filled circles indicate genes with adjusted p-value <0.05 in both contrasts. Hollow circles indicate genes with adjusted p-value >0.05 in one or both contrasts. Green and orange circles = poor- or rich-biased genes that are neither (E) leg- or (F) sex-biased. Dark yellow = rich-biased and L1-biased. Light yellow = nutrition unbiased and L1-biased. Medium yellow = poor-biased and L1- biased. Grey = nutrition unbiased and leg unbiased. Dark brown = rich biased and L3-biased. Light brown = nutrition unbiased and L3-biased. Red = poor-biased and L3-biased. Purple = rich-biased and female-biased. Light pink = nutrition unbiased and female-biased. Dark pink = poor-biased and female-biased. Dark blue = rich-biased and male-biased. Medium blue = nutrition unbiased and male-biased. Turquoise = poor-biased and male-biased. Enrichment of leg- or sex-biased genes in the rear legs of poor and rich males was estimated using the Fisher’s exact tests (**Supplementary Table S2**).

Interestingly, our data also show that nutrition-responsive genes were more frequent in male rear legs (blue bars, **Figure 2A**) compared to male forelegs and to female rear-legs (yellow and purple bars, respectively, **Figure 2A**). The most exaggerated tissue (i.e., the male rear leg under rich treatment) displays the highest number of nutrition-responsive genes with 1595 rich-biased genes (**Figure 2A**, **Table 1**). Surprisingly, the magnitude of nutrition-responsive gene expression was lowest in male rear legs and highest in female rear legs into each nutritional condition (**Figure 2C**). Together, these results show that rich nutrition influences the activity of more genes in male rear legs but with moderate expression levels, whereas the same treatment influences fewer gene but with higher levels in female rear legs. extent in male rear legs. This response becomes dampened with lower nutritional input.

Consistently with this result, we found that leg-biased genes (significantly differentially expressed genes between forelegs and rear legs in males) are two times more numerous in rich than poorly fed males (683 leg-biased genes in rich nutrition versus 308 in poor nutrition, **Table 1**, **Supplementary Figure S6**). However, the magnitude of differential expression for genes with leg-bias expression did not differ between poor and rich treatment (**Supplementary Figure S6**). Moreover, forelegs and rear legs are more similar in absolute length in poorly fed males and more different in rich-fed males. This result therefore suggests that the increase in the degree of leg length divergence in males mobilizes a larger number of differentially expressed genes.

### Rear leg size sexual dimorphism is correlated with higher female-biased gene expression

Rear leg size is highly sexually dimorphic, and the differences between the sexes are either attenuated by poor or exacerbated by rich diet. To test whether the effect of nutrition on leg size dimorphism is mediated through differences in gene expression between the sexes, we compared gene expression profiles of rear legs between males and females reared under rich or poor diet. We found similar numbers of sex-biased genes in poor or rich nutrition with 342 and 319 genes respectively (**Table 1**). Our analysis identified 176 and 247 female-biased genes in individuals raised under poor or rich nutrition, respectively (**Figure 2B**, **Table 1**).

Conversely, we found twice fewer male-biased genes under rich diet in comparison to those fed with poor diet (**Figure 2B**, **Table 1**). Moreover, the magnitude of sex-induced differential expression reflected these differences, with female-biased genes showing the strongest differential expression under rich nutrition and male-biased genes the weakest differential expression under poor nutrition (**Figure 2D**). Thus, increased sexual dimorphism under rich nutrition is associated with an increase in both number and magnitude of female-biased genes, but with a decrease in intensity of male-biased genes. These results suggest that the development of the most pronounced sexual dimorphism observed between male and female rear legs results either from an intense activation in rich females or a strong repression in rich males.

We further examined sex-biased genes that are commonly differentially expressed either in poor or rich nutrition to determine essential genes involved in sexual dimorphism. We found that 53 female-biased genes for only 7 male-biased genes are common in both nutrition treatments (**Supplementary Figure S7**), suggesting that these shared female-biased genes may be involved in a common genetic program associated with sexual dimorphism.

### A specific gene expression signature for male leg exaggeration

Then, we investigated the contribution of sex-biased genes and leg-biased genes to extreme growth variation in response to nutrition. We correlated nutrition-biased genes in male rear legs (poor- or rich-biased genes, y axis in **Figure 2E-F**) with leg-biased genes (L1- or L3- biased genes, x axis in **Figure 2E**) or sex-biased genes (female- or male-biased genes, x axis in **Figure 2F**) in individuals raised under each diets (poor and rich diet are shown in **Figure 2** and **Supplementary Figure S8**, respectively), according to their differential expression level (log2 fold change). We found that rear legs from males reared on rich are enriched in male-biased genes either on poor nutrition (69 out 166 (41%), Fisher’s exact test, p value 5e^-10^, blues dots in **Figure 2E**) or rich nutrition (58 out 72 (80%), Fisher’s exact test, p value 1.15e^-27^, blues dots in **Supplementary Figure S8**). Similarly, male rear legs from rich diet treatment express a high proportion of L3-biased genes either on poor nutrition (55 out 134 (41%), p value 4.1e^-8^, brown dots in **Figure 2E**) or rich nutrition (224 out 361 (62%), p value 3.3e^-69^, brown dot in **Supplementary Figure S8**).

Conversely, when males are reared on poor diet, their rear legs are strongly enriched in female-biased genes (98 out 176 (55%) and 95 out 247 (38%), p value 1.7e-33 and 1.8e-17, pink dots in **Figure 2E** and **Supplementary Figure S8** for female-biased genes on poor and rich condition respectively) and L1-biased genes (66 out 174 (38%) and 106 out 322 (33%), p value 2.8e-12 and 6.5e-14, yellow dots in **Figure 2F**; **Supplementary Figure S8** for L1- biased genes on poor and rich condition respectively). Notably, 77% of the female-biased genes and 54% of the L1-biased genes shared between both nutritional conditions were responsive to nutritional variation in male rear legs. Among these nutrition-responsive genes, 83% of the female-biased genes and 70% of the L1-biased genes were upregulated under poor nutrition, with both categories showing a significantly greater magnitude of differential expression under poor than rich nutrition (**Supplementary Table S4**, **Supplementary Figure S7**). These results suggest that the limited rear leg growth in males fed under poor nutrition is associated with a transcriptomic signature that is highly similar to tissues that do not undergo extreme growth (male forelegs and female rear legs). In contrast, the exaggerated growth of male rear legs under rich nutrition is associated with the activation of a new set of genes that are both male-biased and L3-biased. Together, these results suggest that female-biased and L1-biased genes are associated with a transcriptional state characterized by limited rear-leg growth, whereas male-biased and L3-biased genes are associated with extreme rear-leg growth. Surprisingly, poorly fed male rear legs are also enriched in L3-biased genes found in poor nutrition only (32 out 134 (24%), p value 0.01, red dots in **Figure 2E**) suggesting that despite a limited growth, rear legs in starved males are hyperallometric due to their expression of L3-biased genes.

### The expression of *BMP11* and *Ubx* is nutrition-dependent and L3-biased

We performed a Gene Ontology (GO) enrichment analysis of leg-biased genes in male rear legs under poor or rich nutritional condition separately (**Supplementary Figure S9**). We found that L3-biased genes are enriched in lipid metabolism and cuticle formation processes in both nutrition treatments, with more genes being active under rich nutrition possibly in relation to the high energy costs required to produce an elongated leg especially in rich males. L3-biased genes from rich nutritional treatment contain genes related to carbohydrate metabolic process that are involved in tissue remodeling and energy production, both necessary for growth. Interestingly, rich L3-biased genes also present genes enriched in cell adhesion and development, that are both involved in morphogenetic processes, but also in “multicellular organism development” category, such as *Ultrabithorax* (*Ubx*). By contrast, poor L3-biased genes are enriched in genes involved in proteolysis pathways known to increase during starvation. In addition, only 8% of L3-biased genes are shared between nutritional condition (40 genes are L3-biased in both nutritional conditions, compared to the 321 and 94 L3-biased genes found exclusively in rich and poor nutrition respectively, **Supplementary Figure S10**), including *Ubx* and the growth factor *Bone Morphogenetic Protein 11* (*BMP11*).

### Role of *BMP11* and *Ubx* in leg growth under nutritional influence in *M. longipes*

In our dataset, *Ubx* was surprisingly upregulated in male rear legs under poor compared to rich nutritional treatment. *BMP11* on the other hand was 2,8 times more expressed in rich compared to poor diet. *Ubx* and *BMP11* are known to regulate leg growth during nymphal development in *M. longipes* males (Toubiana et al., 2021b). Since our data show that the expression of both genes is influenced by nutritional treatment, we wanted to test whether these molecular changes link nutritional treatment to differences in leg growth. We therefore performed RNA interference (RNAi) knock-down on juveniles, treated with poor or rich diet, and measured the rear legs and body size of the resulting adult individuals.

*BMP11* knock-down, in males fed on poor diet, induced a 22,7% decrease in the rear legs of males (mean leg length of 3109,9 µm and 4021,3 µm in poor-fed *BMP11* and controls respectively, two-way ANOVA; p < 0.0001; **Figure 3A**, **Supplementary Table S5**) and only 7,5% decrease in rear legs of females (two-way ANOVA; p < 0.01; **Figure 3B**, **Supplementary Table S5**) compared to controls (buffer-injected also fed on poor diet; **Supplementary Table S5**, **Figure 3A-B**). In rich diet, *BMP11* RNAi knockdown induced 29% decrease in rear leg length in males (mean leg length of 3641,8 µm and 5130,6 µm in rich-fed *BMP11* and controls respectively; **Figure 3A**, **Supplementary Table S5**) and 11,5% decrease in female rear legs compared to controls (buffer-injected and fed on rich diet; **Supplementary Table S5**, **Figure 3A-B**). This result suggests that BMP11 knock-down affects more intensely rear leg length in males compared to those in females.

**Figure 3.**
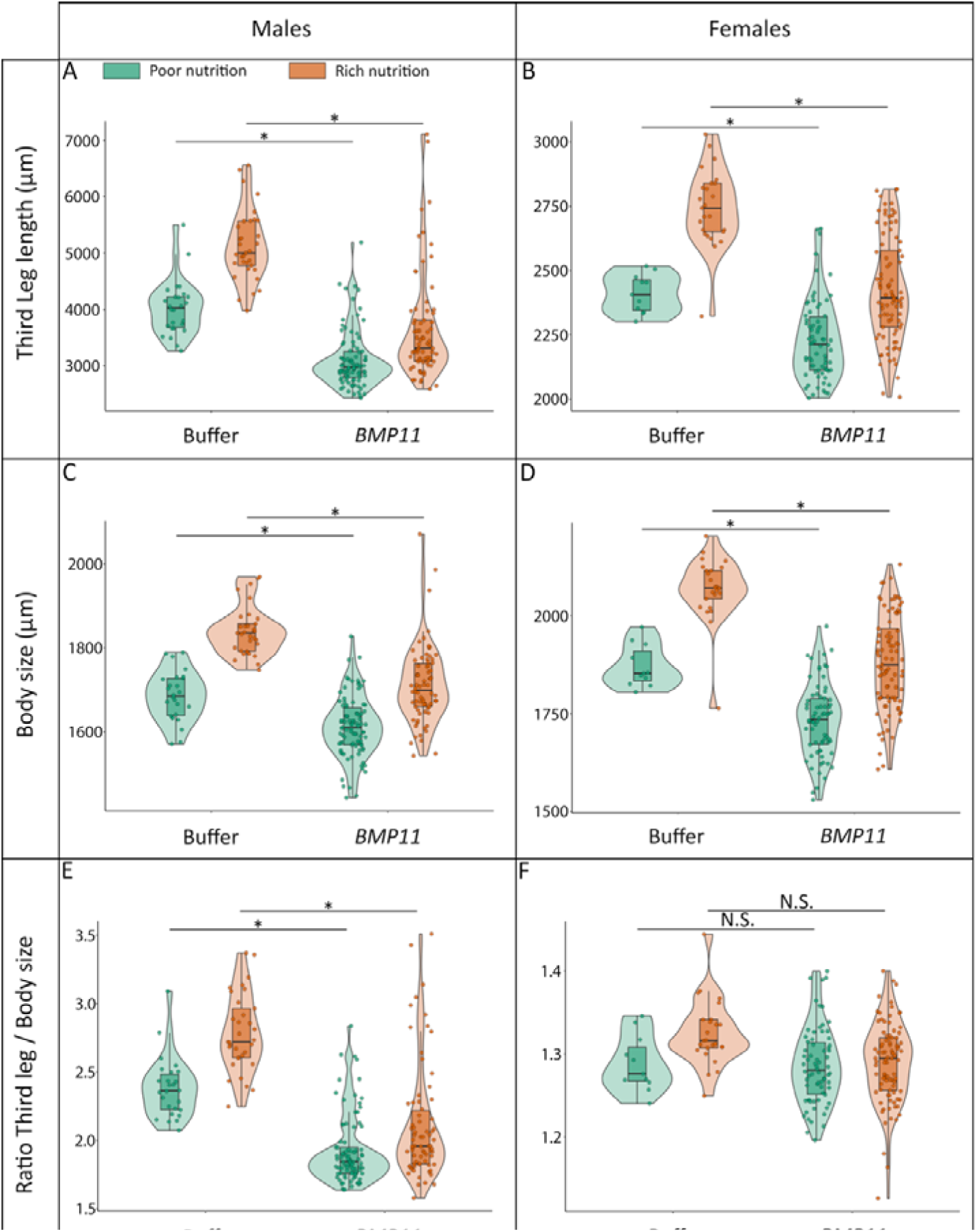
BMP11 knock-down effect on growth of (A-C-E) males and (B-D-F) females in response to nutrition (poor and rich nutrition are represented in green and orange respectively). Measurement are shown for (A, B) rear legs, (C, D) body and (E, F) their relationship in individuals injected either with buffer solution or *bmp11* dsRNA. *BMP11* knock-down induces a reduction in rear leg length and body size in both sexes (*: two-way ANOVA followed by Tukey’s HSD post-hoc tests, p-value <0.05; relative L3 length tested using residuals from L3 length regressed against body size). **G, H** Standardized effect sizes (beta coefficients with 95% confidence intervals) representing the magnitude of (G) nutritional and (H) knock-down treatment. The effect size is represented in males and females (squares and circles, respectively). (G) Nutritional effect sizes are shown for individuals injected with buffer or *BMP11* dsRNA (blue and pink, respectively). (H) Knock-down effect sizes are shown for individuals reared under poor or rich nutritional conditions (green and orange, respectively). Differences in effect sizes between treatments were tested using the interaction term of the linear model (* : p-value <0.05).

Moreover, the effect of *BMP11* knock-down on male rear leg length and body size depended on nutritional condition, as indicated by a significant treatment x nutrition interaction (t = - 2,74 and -2,44, p-value = 0,0067 and 0,016 for male rear leg length and body size, respectively). Accordingly, the standardized knock-down effect size was 38,8% stronger under rich nutrition than poor nutrition in male rear legs (ß = 0,95, 95% CI [0,63 ; 1,26] and ß = 1,55, 95% CI [1,25 ; 1,84], respectively, **Figure 3H**) and 42,9% stronger in rich-fed male body (ß = 0.67, 95% CI [0,36 ; 0,98] and 1.20, 95% CI [0,90 ; 1,49] in poor and rich nutrition, respectively, **Figure 3H**). Contrariwise, the treatment x nutrition interaction was not significant in females neither in rear leg length (t = - 1,95, p = 0,053) or body size (t = -1,22, p = 0,224). Together, these results suggest that the effect of *BMP11* on male leg growth is more pronounced in rich relative to poor diet, resulting in similar rear leg length distribution in *BMP11* males fed under poor and rich diet. Conversely, the same interaction can be expressed in terms of nutritional sensitivity: the standardized effect of nutrition in male rear leg length is reduced by 52% in *BMP11* compared to control males (ß = 0,55, 95% CI [0,33 ; 0,77] and 1,15, 95% CI [0,78 ; 1,52], respectively) and by 36% in *BMP11* male body (ß = 0,91, 95% CI [0,69 ; 1,13] and 1,44, 95% CI [1,07 ; 1,80] in *BMP11* and control males, respectively). The stronger effect of *BMP11* knock-down on male rear leg length was supported after accounting for variation in body size. *BMP11* knock-down significantly reduced body size-corrected rear leg length in males under poor and rich nutrition (two-way ANOVA; p < 0.0001; **Figure 3E**, **Supplementary Table S5**) indicating that the reduction in rear leg length cannot be explained only by the reduction in overall body size. In contrast, *BMP11* knock-down did not significantly affect body size-corrected rear leg length in females (two-way ANOVA; p > 0.05; **Figure 3F**, **Supplementary Table S5**). Together, these results show that BMP11 knock-down disproportionately reduces rear leg growth under rich nutrition specifically in males, thereby reducing nutritional plasticity of the exaggerated male rear legs.

We conclude that BMP11 is required for nutritional plasticity during the development of exaggerated rear legs in male.

Ubx has been shown to regulate leg growth in water striders in a dose-dependent manner, with high levels of Ubx having a negative effect on leg length [34]. We therefore hypothesized that the over-expression of *Ubx* seen in poor diet treatment acts to slow down the growth of the rear legs of *M. longipes* males. To test this hypothesis, we injected two different concentrations of double-stranded RNA (dsRNA), 0,5 and 5 µg/µL, of *Ubx* in an attempt to manipulate the levels of remaining *Ubx* mRNA in leg tissues, with lower concentration of *Ubx* dsRNA leading to moderate reduction of Ubx levels [34]. However, we were not able to see any significant differences neither in rear legs length or body size in males between the two concentrations of injected dsRNA within each nutrition (two-way ANOVA, p-value >0.05, **Supplementary Figure S11**). This experiment showed that regardless of *Ubx* dsRNA concentration, *Ubx* RNAi results in shorter male rear legs by both 18% or by 14% when insects were reared on poor or rich diet (**Supplementary Table S6**, **Figure 4A**). Body size in these males was unaffected by *Ubx* knockdown, suggesting that its role during nymphal development is restricted to the legs (**Figure 4C**). Moreover, *Ubx* knock-down does not show nutritional effect size variation in rear leg length since treatment x nutrition interaction fails significance tests (t = -0,287, p-value = 0,775), indicating that *Ubx* is not involved in the nutrition-dependent plasticity. In females, however, the effect of *Ubx* knock-down on absolute rear leg length differed between nutritional condition (t = -2.914, p-value = 0.0039). The standardized knock-down effect was stronger in female rear legs under poor nutrition than rich nutrition (ß = 1,19, 95% CI [0,99; 1,40] and ß = 0,71, 95% CI [0,46; 0,96], respectively) corresponding to reduction in rear legs by approximatively 10% and 5%, respectively (**Figure 4B**, **Supplementary Table S6**). A significant treatment x nutrition interaction was also detected for female body size (t = -2.436, p = 0.0156). However, the magnitude of this effect was small, with *Ubx* knock-down reducing body size by only approximately 1.8% under poor nutrition (from 1933 µm in controls to 1898 µm in *Ubx* females; **Figure 4D**; **Supplementary Table S6**). Despite these interactions for absolute rear-leg length and body size, nutrition did not significantly affect body size-corrected rear leg length in *Ubx* females (*Ubx* Poor vs *Ubx* Rich, p > 0.05). Thus, although *Ubx* knock-down produced a statistically detectable nutrition-dependent effect on absolute rear leg length in females, this effect was modest and was not associated with a detectable change in rear-leg length relative to body size. Together, these results fail to implicate Ubx to the nutrition-dependent plasticity of the exaggerated male rear legs in *Microvelia longipes*.

**Figure 4.**
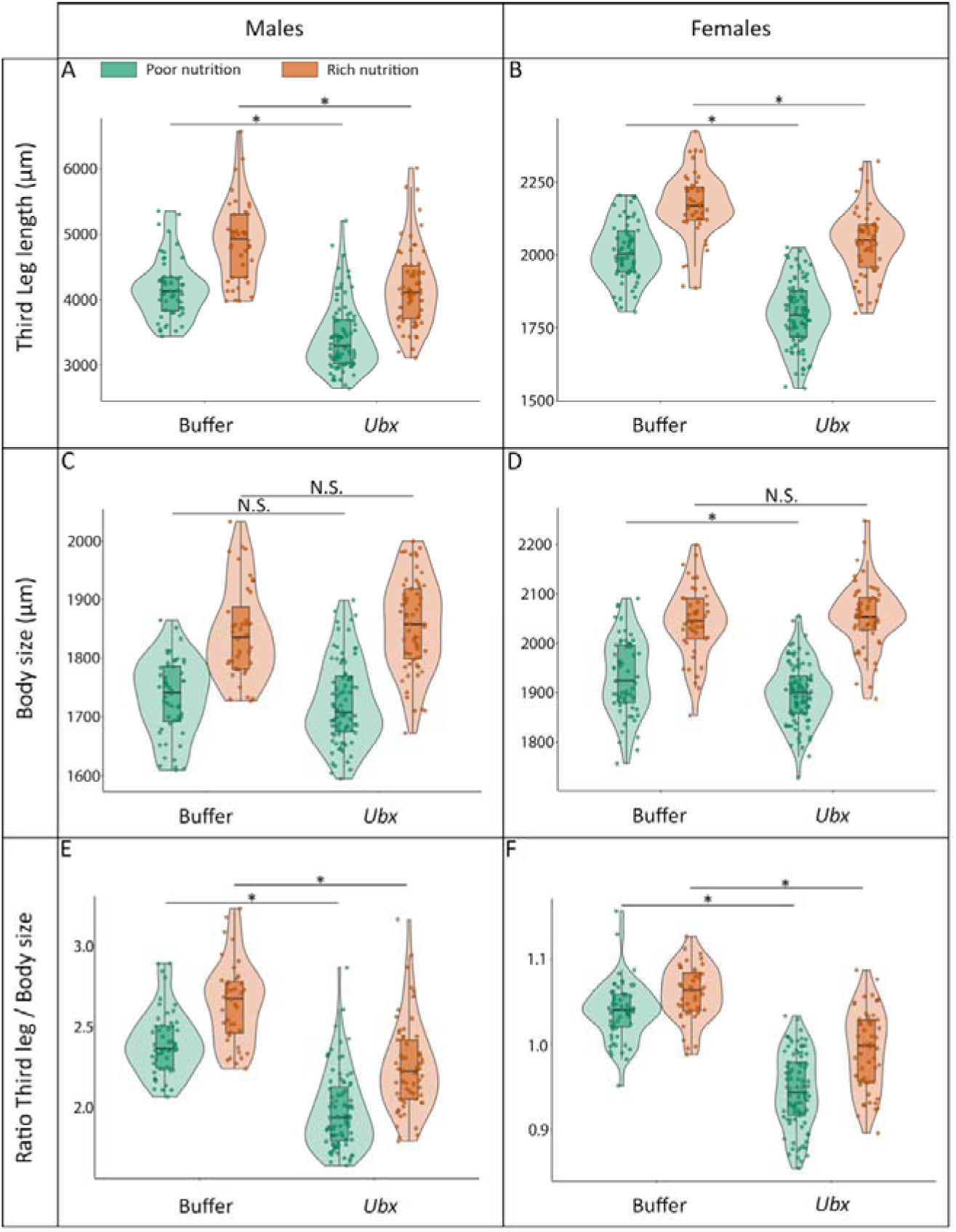
Ubx knock-down effect on growth of (A-C-E) males and (B-D-F) females in response to nutrition (poor and rich nutrition are represented in green and orange respectively). Measurement are shown for (A, B) rear legs, (C, D) body and (E, F) their relationship in individuals injected either with buffer solution or *ubx* dsRNA. *Ubx* knock-down induces a reduction in rear leg length in both sexes but has no effect in body size, expected the small but significant reduction in body size in poor females (* : two way ANOVA followed by Tukey’s HSD post-hoc tests, p-value <0.05; relative L3 length tested using residuals from L3 length regressed against body size). **G, H** Standardized effect sizes (beta coefficients with 95% confidence intervals) representing the magnitude of (G) nutritional and (H) knock-down treatment. The effect size is represented in males and females (squares and circles, respectively). (G) Nutritional effect sizes are shown for individuals injected with buffer or *Ubx* dsRNA (blue and pink, respectively). (H) Knock-down effect sizes are shown for individuals reared under poor or rich nutritional conditions (green and orange, respectively). Differences in effect sizes between treatments were tested using the interaction term of the linear model (* : p-value <0.05).

## Discussion

Our study provides insights into the molecular processes linking nutritional input to the development of an exaggerated sexually selected trait—a central yet understudied question in evolutionary biology. Using the water strider *Microvelia longipes* as a model, we demonstrate that nutritional variation drives extreme phenotypic plasticity in male rear leg length, a trait critical for male-male competition and reproductive success [21]. We reveal how environmental cues are translated into molecular and phenotypic variation, offering novel insights into the proximal mechanisms underlying the development of exaggerated sexually selected traits.

### Nutritional plasticity as a primary modulator of gene expression and trait development

Our study in *Microvelia longipes* provides critical insights into the molecular processes linking nutritional input to the development of an exaggerated sexually selected trait. Using reaction norm experiments, transcriptomics, and functional validation by RNAi, we demonstrate that nutritional variation drives extreme phenotypic plasticity in male rear leg length, a trait essential for male-male competition and reproductive success [21]. Our results show that nutrition is the primary driver of gene expression variation in *M. longipes* legs, surpassing the effects of serial homology (leg type) or sex. This aligns with studies in other insects, including those where trait exaggeration is carried by females [35, 36]. However, other studies in beetles came to different conclusions. For example, Kijomoto et al; found that gene expression varies the most between tissues and sex than between nutritional treatments [37]. This contrasting conclusion could be the result of comparing non-homologous tissues between the sexes (horn vs epidermis) and possible due to the use of genetically diverse samples [37]. In *M. longipes*, male rear legs, which undergo the most pronounced nutrition-dependent growth, exhibit the strongest transcriptional response to diet, characterized by a higher number of nutrition-responsive genes rather than larger changes in the magnitude of expression. This suggests that a broad, fine-tuned transcriptional response underlies the sensitivity of male rear legs to nutritional variation, consistent with the “many small effects” model of complex trait evolution [38].

While nutrition is known to shape phenotypic plasticity in secondary sexual traits [7, 9], previous studies have linked the degree of exaggeration and nutritional responsiveness to changes in gene expression under dietary input [35, 36], often involving the Insulin pathway [15, 18]. However, in *M. longipes*, neither this nor prior studies [22] have linked leg length exaggeration to Insulin signaling, as RNAi knockdown of Insulin Receptors did not alter male leg growth. Instead, other genes, such as BMP11, play a functional role in mediating nutritional plasticity.

### Sexual dimorphism and the transcriptomic signature of exaggeration

Sexual dimorphism in *M. longipes* rear legs is amplified under rich nutrition, with males developing disproportionately longer legs than females. Our data reveal that this pronounced dimorphism correlates with higher number of female-biased genes under rich diet, while male-biased genes are less frequent. This suggests that strong sexual dimorphism in leg length may be associated with a shift toward female-biased gene expression, rather than with an increased contribution of male-biased genes. This shift could result from either increased expression in females or decreased expression in males. This aligns with studies in *Drosophila* where sex-specific traits are often regulated by repression of the opposite sex’s developmental pathways [39].

The enrichment of male-biased and L3-biased (rear leg-specific) genes in rich-fed males points to a transcriptomic signature unique to exaggerated growth. Conversely, poor-fed males show enrichment for female-biased and L1-biased (foreleg-like) genes, suggesting that limited rear leg growth in starved males resembles the transcriptional state of non-exaggerated tissues. This mirrors findings in the beetle *Onthophagus taurus*, where *doublesex* and FoxO mediate sex- and nutrition-specific growth programs [16, 18]. Our results extend this framework by showing that continuous traits may rely on similar molecular players but with graded, rather than switch-like, responses to nutrition.

### Functional contribution of BMP11 and Ubx to nutrition-dependent extreme growth

The contrasting functional effects of BMP11 and Ubx highlight an important distinction. Nutritional responsiveness in gene expression does not necessarily imply a functional role in nutritional plasticity. BMP11 expression increases under rich nutrition in male rear legs, and its contribution to rear leg growth is stronger under rich diet condition. This suggests that BMP11 mediates the conversion of higher nutritional intake into growth, consistent with the resource allocation hypothesis [40]. Accordingly, BMP11 knockdown substantially reduces the nutritional sensitivity of male rear leg growth, even after accounting for body size variation, supporting its specific role in coupling nutrition to extreme growth.

In contrast, Ubx knockdown reduces rear leg length equally under both nutritional conditions, indicating that while Ubx is required for rear leg growth, it does not mediate its nutrition-dependent plasticity. This aligns with findings in *Gerris* water striders, where Ubx inhibits leg growth when Ubx dose is high [34], but its role appears independent of nutritional state in *M. longipes*. Thus, BMP11 and Ubx contribute to distinct aspects of plasticity; BMP11 mediates nutritional responsiveness, while Ubx regulates basal growth patterns (**Figure 5**).

**Figure 5.**
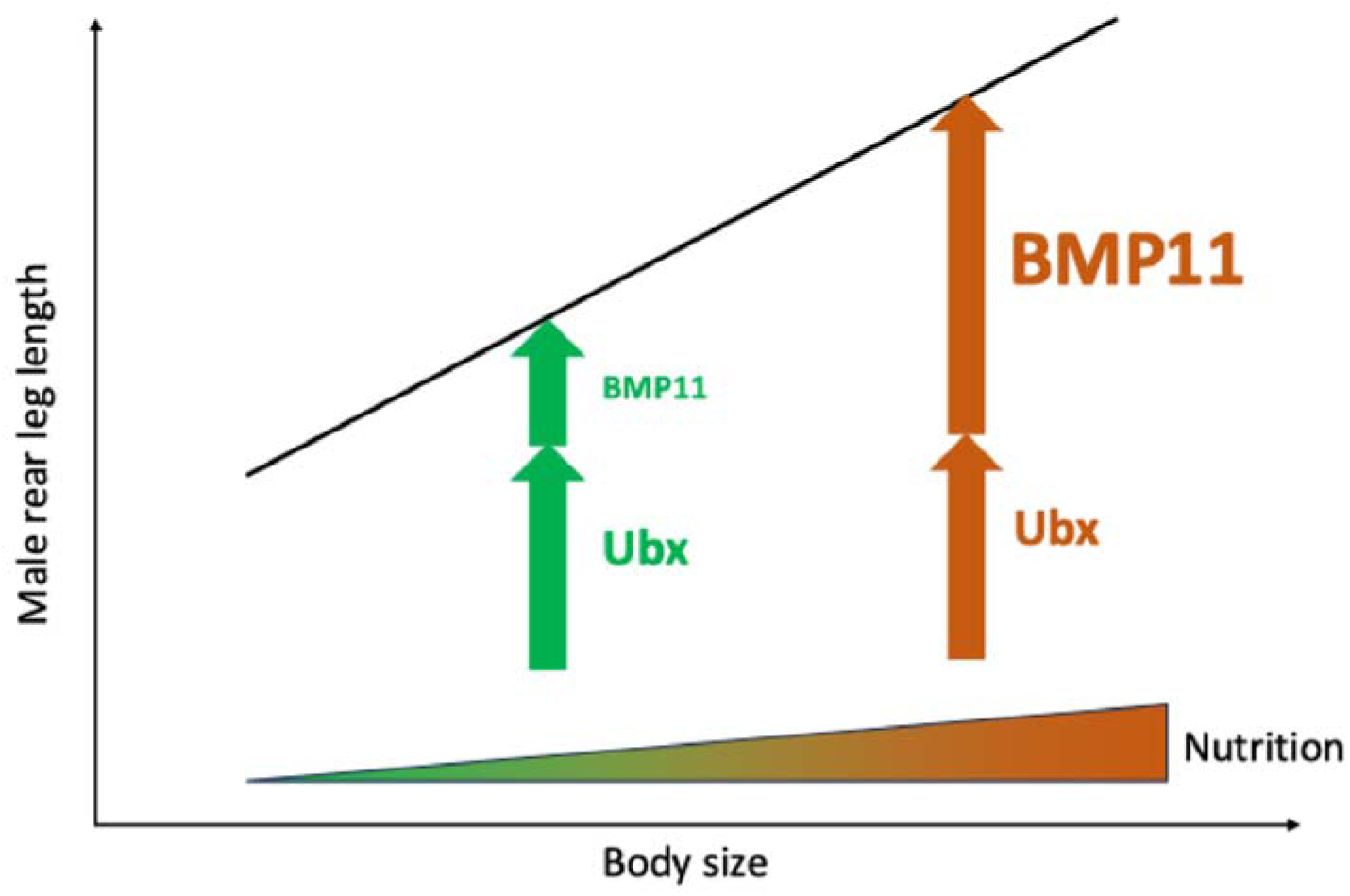
Graphical summary: Mechanisms underlying nutrition-dependent extreme growth variation in the rear legs of *Microvelia longipes* males. Ubx promotes rear-leg elongation with the same magnitude under both nutritional conditions, indicating Ubx is not involved in nutrition-dependent growth variation. In contrast, BMP11 strongly promotes rear leg length in rich-fed male third legs, and moderately in poor-fed males. As BMP11 is upregulated in rear legs of well-fed individuals, its nutrition-dependent expression and function make BMP11 a key mediator of nutritional plasticity in exaggerated male rear legs.

Interestingly, BMP11 is involved in extreme growth in a sex-specific manner. Although BMP11 expression does not differ significantly between males and females, its knockdown has a stronger effect on leg growth in male rear legs, suggesting that its contribution to exaggerated growth depends on both its expression level and the sex in which it acts. This is consistent with studies showing that gene expression can be differentially altered by environmental cues in males and females [41].

Our data also suggest that the development of exaggerated male rear legs involves a shift between transcriptional programs associated with non-exaggerated and exaggerated states. Poor-fed male rear legs retain a transcriptional signature similar to non-exaggerated tissues, with an enrichment of female- and L1-biased genes. Conversely, rich-fed males show enrichment for male- and L3-biased genes, indicating that extreme growth is associated with the activation of a male- and rear leg-specific transcriptional program. This dual mechanism consisting of repression of genes associated with the non-exaggerated state and activation of gene that promote growth, may underlie the nutrition-dependent modulation of trait exaggeration in a sex specific manner.

### Plasticity as a driver of evolutionary innovation

Most studies on exaggerated traits focus on holometabolous insects with discrete morphs [9, 19, 42]. In contrast, *M. longipes*, a hemimetabolous insect with continuous male leg length variation, relies on graded expression of genes like BMP11 to generate plasticity. This suggests that while conserved genetic toolkits (e.g., BMP, Ubx) underlie exaggerated traits across taxa, their regulatory architecture (discrete vs. continuous) shapes the form of plasticity. Future work could explore whether hemimetabolous insects generally rely on graded transcriptional responses, while holometabolous insects favor threshold-based mechanisms. Phenotypic plasticity is increasingly recognized as a catalyst for evolutionary change [3, 5]. Our findings in *M. longipes* support this view by showing that nutritional plasticity in leg growth is mediated by genes (BMP11, Ubx) with deep developmental roles. The decoupling of genetic and environmental effects, where identical genotypes produce divergent phenotypes under different diets—highlights how plasticity can unmask cryptic genetic variation [2], facilitating the evolution of novel, exaggerated traits.

### Future directions

While our study identifies BMP11 as a key regulator of nutritional plasticity, the upstream signals (e.g., insulin, ecdysone, or TOR pathways) linking nutrition to their expression remain unknown, despite their established roles in growth regulation in other insects [43–45].

Additionally, epigenetic mechanisms, such as DNA methylation or histone modifications (Richards, 2006), may contribute to the stable, nutrition-dependent gene expression patterns we observed but were not explored here. Our use of inbred lines controlled for genetic background, but genotype-by-environment interactions in natural populations could further modulate plasticity [46, 47]. Finally, while RNAi knockdowns provided functional insights, CRISPR-based knockouts or tissue-specific manipulations could offer more precise validation of the roles of BMP11 and Ubx.

## Conclusion

In *Microvelia longipes*, nutritional input and gene expression interact to produce one of the most extreme examples of phenotypic plasticity in nature. Our work demonstrates that the modulation of transcriptional programs underlies the effects of nutrition on the development of variable exaggerated traits, with BMP11 acting as a central component in this regulation. These findings advance our understanding of how environmental factors shape complex phenotypes and provide a molecular entry point for studying the evolution of plasticity under sexual selection. By illuminating the proximate mechanisms behind trait exaggeration, we pave the way for future research into the ultimate causes of diversity in nature.

## Supporting information

SOM

## Authors’ contributions

A.K., K.P. and I.D. designed research; I.D. performed experiments, RNA-seq analysis; S.V. and I.D. did the measurements, I.D. analyzed data; and A.K. and I.D. wrote the paper.

## Acknowledgements

We thank Juliette Mendes for help with insect care and the Khila lab members for discussions. Sequencing was performed by the SNP&SEQ Technology Platform in Uppsala and IGFL Plateforme de Sequençage. The facility is part of the National Genomics Infrastructure (NGI) Sweden and Science for Life Laboratory. The SNP&SEQ Platform is also supported by the Swedish Research Council and the Knut and Alice Wallenberg Foundation. We thank the IGFL Plateforme de Sequençage, Benjamin Gillet, Sandrine Hughes for help with sequencing. We gratefully acknowledge support from the CBPsmn (PSMN, Pôle Scientifique de Modélisation Numérique) of the ENS de Lyon for the computing resources. The platform operates the SIDUS solution developed by Emmanuel Quemener (https://www.linuxjournal.com/content/sidus%E2%80%94-solution-extreme-deduplication-operating-system). This work was supported by an FRM-équipe grant EQU202103012573 to AK.

## References

1. Darwin, C., On the origin of species by means of natural selection, or, The preservation of favoured races in the struggle for life. 1859, London: J. Murray. ix, [1], 502, 32, [1] fold. leaf of plates (32 at end advertisements).

2. Beldade, P., A.R. Mateus, and R.A. Keller, Evolution and molecular mechanisms of adaptive developmental plasticity. Mol Ecol, 2011. 20(7): p. 1347–63.

3. Gilbert, S.F. and D. Epel, Ecological Developmental Biology. 2009, Sunderland: Sinauer Associates, Inc.

4. Schlotterer, C., et al., Combining experimental evolution with next-generation sequencing: a powerful tool to study adaptation from standing genetic variation. Heredity (Edinb), 2015. 114(5): p. 431–40.

5. West-Eberhard, M.J., Developmental plasticity and the origin of species differences. Proc Natl Acad Sci U S A, 2005. 102 Suppl 1: p. 6543–9.

6. Darwin, C., The descent of man, and selection in relation to sex. 1871, London,: J. Murray. 2 v.

7. Bonduriansky, R., The evolution of condition-dependent sexual dimorphism. The American naturalist, 2007. 169(1): p. 9–19.

8. Bonduriansky, R., Condition dependence of developmental stability in the sexually dimorphic fly Telostylinus angusticollis (Diptera: Neriidae). J Evol Biol, 2009. 22(4): p. 861–72.

9. Emlen, D.J. and H.F. Nijhout, The development and evolution of exaggerated morphologies in insects. Annual Review of Entomology, 2000. 45: p. 661–708.

10. Shingleton, A.W., The regulation of organ size in Drosophila: physiology, plasticity, patterning and physical force. Organogenesis, 2010. 6(2): p. 76–87.

11. Shingleton, A.W. and W.A. Frankino, New perspectives on the evolution of exaggerated traits. Bioessays, 2013. 35(2): p. 100–7.

12. Gilbert, S.F., Mechanisms for the environmental regulation of gene expression: ecological aspects of animal development. J Biosci, 2005. 30(1): p. 65–74.

13. Grath, S. and J. Parsch, Sex-Biased Gene Expression. Annual Review of Genetics, Vol 50, 2016. 50: p. 29–44.

14. Mank, J.E., The transcriptional architecture of phenotypic dimorphism. Nature Ecology & Evolution, 2017. 1(1).

15. Emlen, D.J., et al., A mechanism of extreme growth and reliable signaling in sexually selected ornaments and weapons. Science, 2012. 337(6096): p. 860–4.

16. Kijimoto, T., A.P. Moczek, and J. Andrews, Diversification of doublesex function underlies morph-, sex-, and species-specific development of beetle horns. Proceedings of the National Academy of Sciences of the United States of America, 2012. 109(50): p. 20526–20531.

17. Lavine, L., et al., Exaggerated Trait Growth in Insects. Annual Review of Entomology, Vol 60, 2015. 60: p. 453–472.

18. Casasa, S. and A.P. Moczek, Insulin signalling’s role in mediating tissue-specific nutritional plasticity and robustness in the horn-polyphenic beetle. Proceedings of the Royal Society B-Biological Sciences, 2018. 285(1893).

19. Moczek, A.P., C.A. Brühl, and F.T. Krell, Linear and threshold-dependent expression of secondary sexual traits in the same individual: insights from a horned beetle (Coleoptera: Scarabaeidae). Biological Journal of the Linnean Society, 2004. 83(4): p. 473–480.

20. Andersen, N.M., The semiaquatic bugs (Hemiptera: Gerromorpha). Vol. - Entomonograph Vol. 3. . 1982, Klampenborg, Denmark.: Scandinavian Science Press LTD.

21. Toubiana, W. and A. Khila, Fluctuating selection strength and intense male competition underlie variation and exaggeration of a water strider’s male weapon. Proc Biol Sci, 2019. 286(1901): p. 20182400.

22. Toubiana, W., et al., The growth factor BMP11 is required for the development and evolution of a male exaggerated weapon and its associated fighting behavior in a water strider. Plos Biology, 2021. 19(5).

23. Pruvot, C., et al., Sexual conflict, directional sexual selection and phenotypic plasticity jointly drive the evolution of extreme phenotypic variation. bioRxiv, 2026: p. 2026.08.18.745420.

24. Bolger, A.M., M. Lohse, and B. Usadel, Trimmomatic: a flexible trimmer for Illumina sequence data. Bioinformatics, 2014. 30(15): p. 2114–2120.

25. Ewels, P., et al., MultiQC: summarize analysis results for multiple tools and samples in a single report. Bioinformatics, 2016. 32(19): p. 3047–8.

26. Kopylova, E., L. Noé, and H. Touzet, SortMeRNA: fast and accurate filtering of ribosomal RNAs in metatranscriptomic data. Bioinformatics, 2012. 28(24): p. 3211–3217.

27. Danecek, P., et al., Twelve years of SAMtools and BCFtools. Gigascience, 2021. 10(2).

28. Liao, Y., G.K. Smyth, and W. Shi, featureCounts: an efficient general purpose program for assigning sequence reads to genomic features. Bioinformatics, 2014. 30(7): p. 923–930.

29. Pertea, M., et al., Transcript-level expression analysis of RNA-seq experiments with HISAT, StringTie and Ballgown. Nat Protoc, 2016. 11(9): p. 1650–67.

30. Toubiana, W., et al., Impact of male trait exaggeration on sex-biased gene expression and genome architecture in a water strider. Bmc Biology, 2021. 19(1).

31. Love, M.I., W. Huber, and S. Anders, Moderated estimation of fold change and dispersion for RNA-seq data with DESeq2. Genome Biol, 2014. 15(12): p. 550.

32. Khila, A., E. Abouheif, and L. Rowe, Comparative Functional Analyses of Ultrabithorax Reveal Multiple Steps and Paths to Diversification of Legs in the Adaptive Radiation of Semi-Aquatic Insects. Evolution, 2014.

33. Finet, C., et al., The achaete-scute complex contains a single gene that controls bristle development in the semi-aquatic bugs. Proceedings of the Royal Society B-Biological Sciences, 2018. 285(1892).

34. Refki, P.N., et al., Emergence of tissue sensitivity to Hox protein levels underlies the evolution of an adaptive morphological trait. Developmental Biology, 2014. 392(2): p. 441–453.

35. Casasa, S., E.E. Zattara, and A.P. Moczek, Nutrition-responsive gene expression and the developmental evolution of insect polyphenism. Nature Ecology & Evolution, 2020. 4(7): p. 970-+.

36. Qi, C.C., W.Q. Zhang, and Y.G. Hu, Transcriptomic basis underlying the evolution of female horn plasticity in scarab beetles. Frontiers in Zoology, 2026. 23(1).

37. Kijimoto, T., et al., The nutritionally responsive transcriptome of the polyphenic beetle Onthophagus taurus and the importance of sexual dimorphism and body region. Proceedings of the Royal Society B-Biological Sciences, 2014. 281(1797).

38. Rockman, M.V., The QTN program and the alleles that matter for evolution: all that’s gold does not glitter. Evolution, 2012. 66(1): p. 1–17.

39. Williams, T.M., et al., The regulation and evolution of a genetic switch controlling sexually dimorphic traits in Drosophila. Cell, 2008. 134(4): p. 610–23.

40. Bonduriansky, R., Sexual selection and allometry: a critical reappraisal of the evidence and ideas. Evolution; international journal of organic evolution, 2007. 61(4): p. 838–49.

41. Camus, M.F., A. Chakraborty, and M. Reuter, Disentangling the reproductive and metabolic transcriptional responses to diet in Drosophila melanogaster. G3-Genes Genomes Genetics, 2026. 16(4).

42. Toubiana, W. and A. Khila, The benefits of expanding studies of trait exaggeration to hemimetabolous insects and beyond morphology. Current Opinion in Genetics & Development, 2016. 39: p. 14–20.

43. Colombani, J., et al., Antagonistic actions of ecdysone and insulins determine final size in Drosophila. Science, 2005. 310(5748): p. 667–670.

44. Colombani, J., et al., A nutrient sensor mechanism controls growth. Cell, 2003. 114(6): p. 739–749.

45. Layalle, S., N. Arquier, and P. Léopold, The TOR Pathway Couples Nutrition and Developmental Timing in. Developmental Cell, 2008. 15(4): p. 568–577.

46. Via, S., et al., Adaptive Phenotypic Plasticity - Consensus and Controversy. Trends in Ecology & Evolution, 1995. 10(5): p. 212–217.

47. Via, S. and R. Lande, Genotype-Environment Interaction and the Evolution of Phenotypic Plasticity. Evolution, 1985. 39(3): p. 505–522.

