## Supplementary material for "Nutrition mediates extreme growth variation through deep changes in gene expression in the water strider *Microvelia longipes*": SOM

**Author affiliations:**

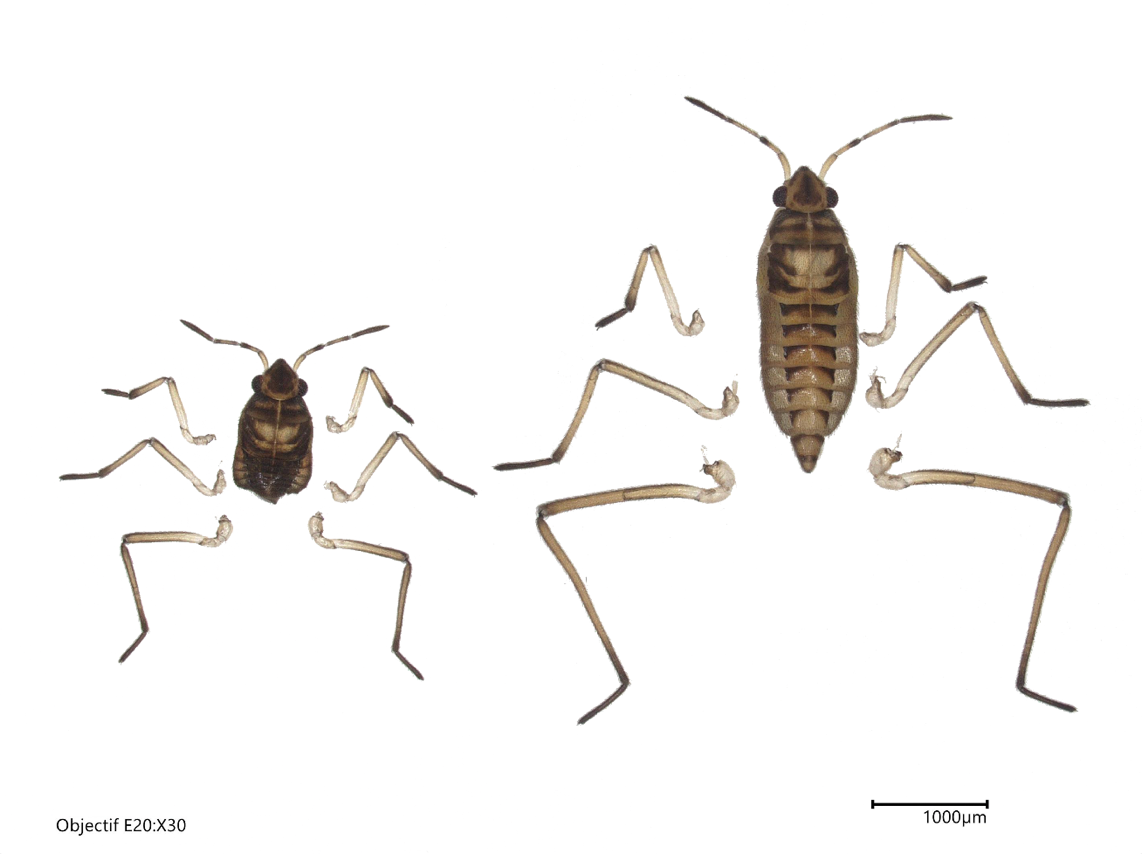
**Supplementary figures:**

**Supplementary figure S1:** Example of two extremes phenotypes in last nymphal instar males developed under poor (left) or rich (right) nutrition. These males belong of the limit of growth range and illustrate how the difference in length is visible to naked eyes.

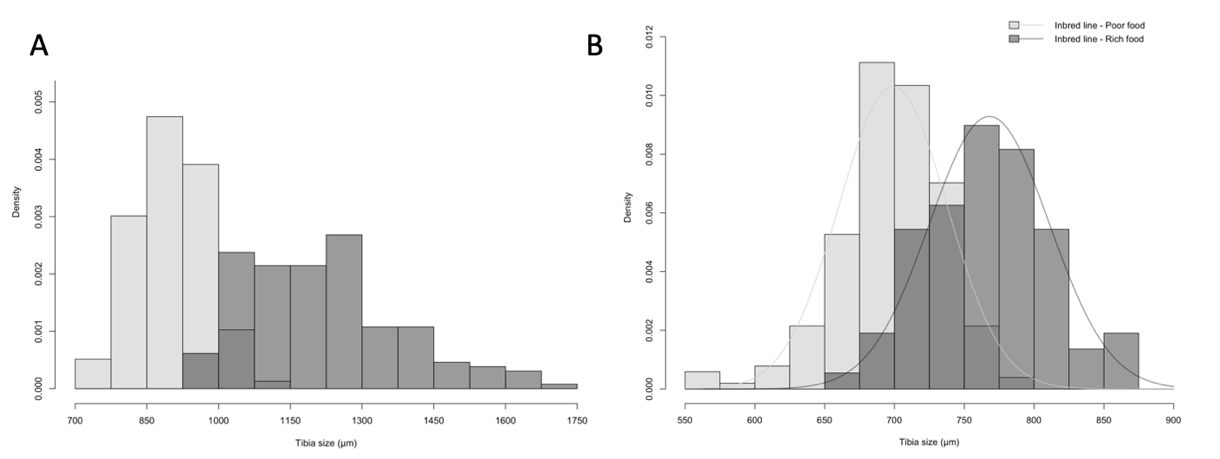

**Supplementary figure S2:** Variation in rear tibia length in (a) males and (b) females selected for RNA-seq. The nymphs have been raised under rich (dark grey) or poor diet (light grey) until the 5^th^ nymphal instar where they were collected for RNA extraction. Individuals from the 3 RNA-seq biological replicates are pooled in this figure. After the selection biased on absolute rear leg size, the variation in length between condition increase in males (∆=315µm,45 µm, W = 645.5, p-value < 2.2e-16), while is always small in females (∆=69,04 µm, W = 3378, p-value < 2.2e-16). Selected males are significantly bigger or smaller than the non-selected males (Wilcoxon test, p = 2.2e-08 and p = 2.3e-19 in poor and rich diet respectively)

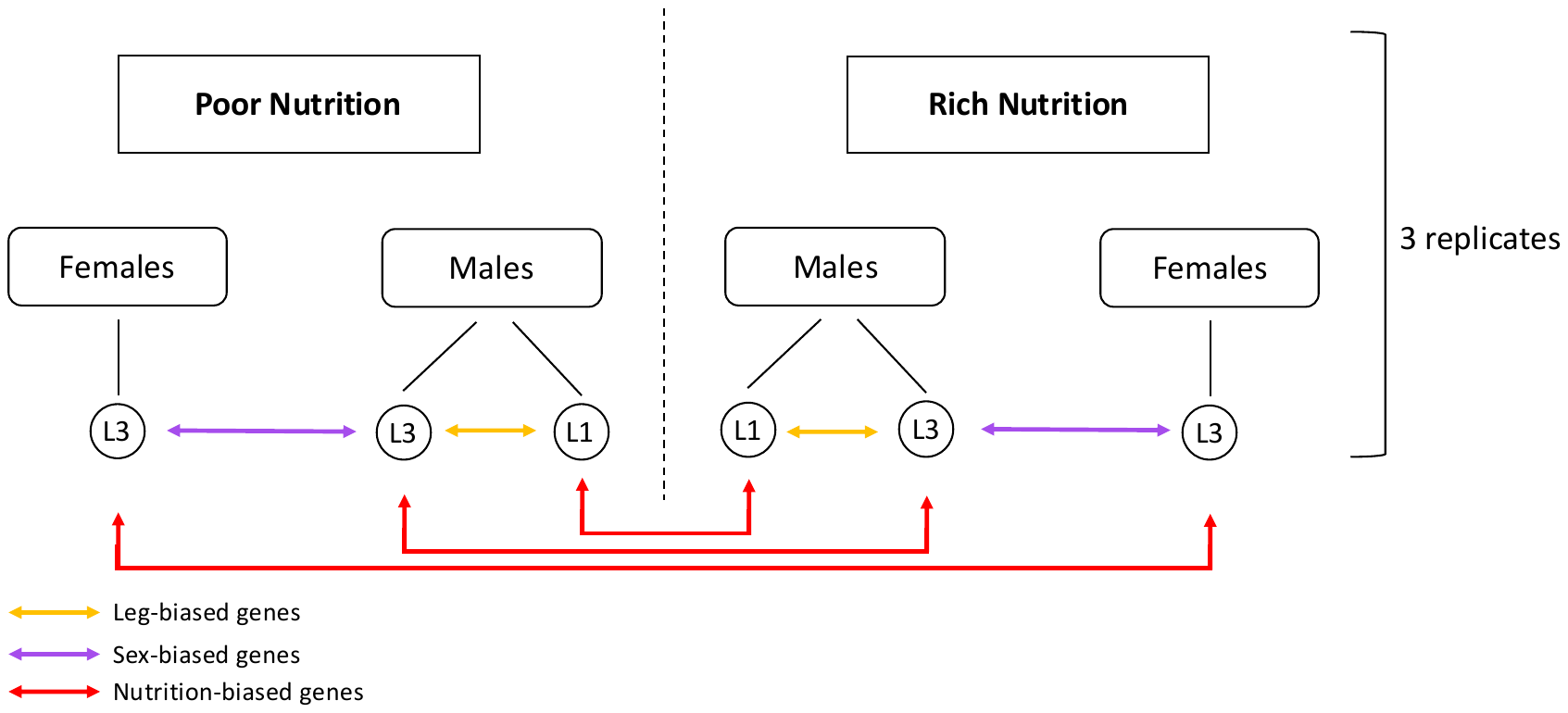

**Supplementary figure S3:** Experimental design of the comparative transcriptomic analysis. It represents the three different comparisons used in the transcriptomic analysis, namely the nutrition, sex and leg condition. This comparative analysis was performed on three replicates.

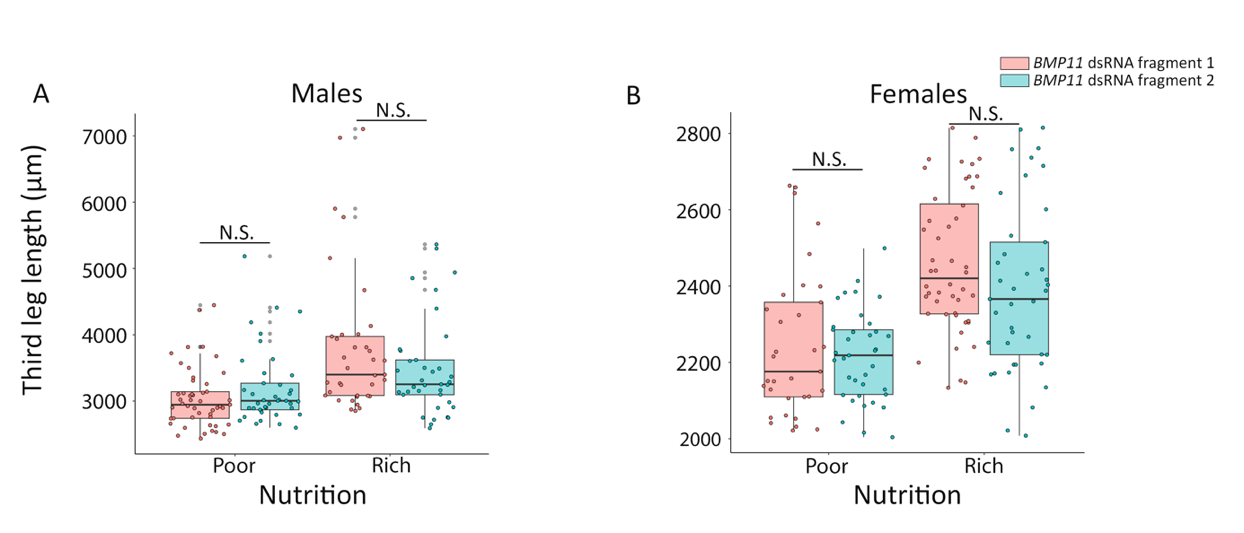

**Supplementary figure S4:** Rear leg length in adults (**A**) males and (**B**) females injected with dsRNA against two non-overlapping *BMP11* fragments, namely fragment 1 and fragment 2 (pink and blue, respectively). Males were raised under poor or rich nutrition after dsRNA injection. There are no significant differences in length between *BMP11* fragments neither in poor or rich nutrition (p-value > 0.05, two-way ANOVA followed by EMM pairwise comparisons within each nutritional condition).

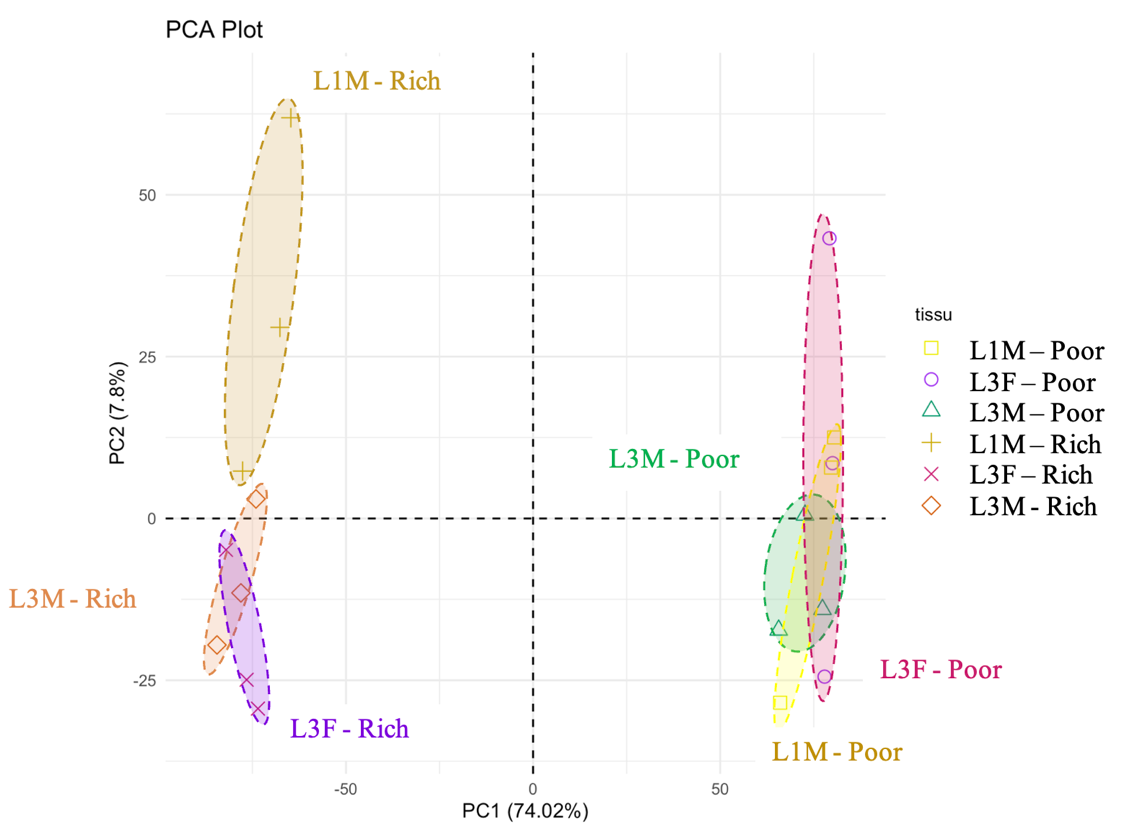

**Supplementary figure S5**: Principal component analysis (PCA) on the whole transcriptomic data of male first and third legs and female third legs raised under poor or rich nutrition. **c** First axis (PC1) explains differences between nutritional treatments while PC2 explains differences between leg type.

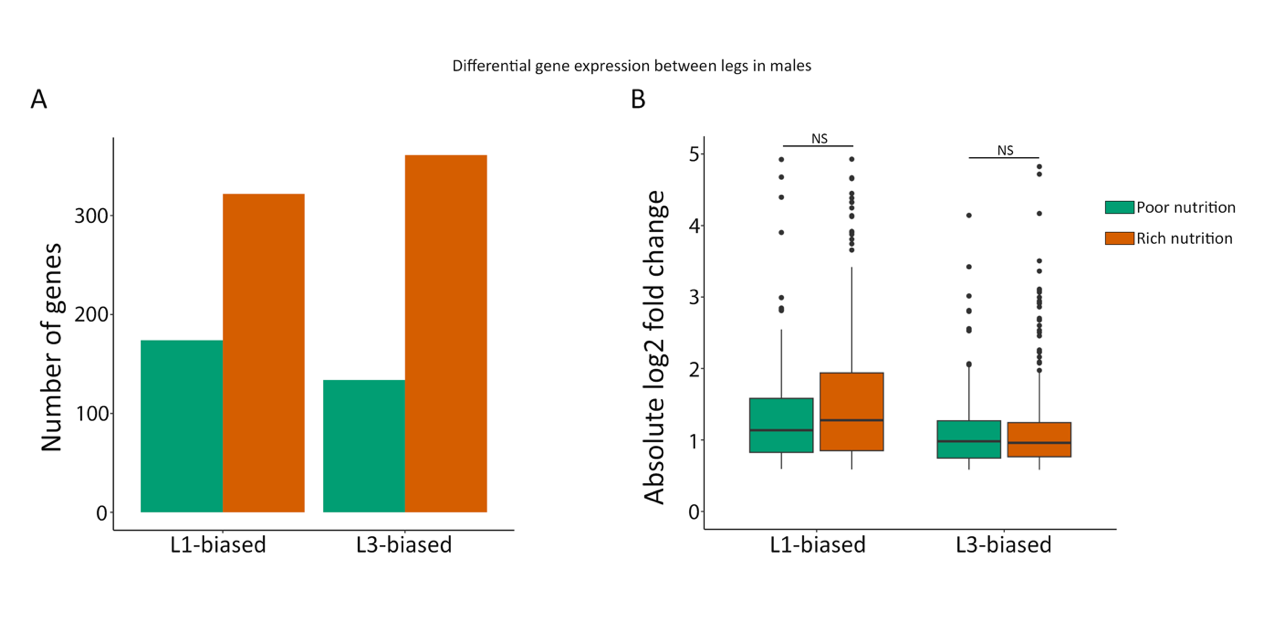

**Supplementary figure S6**: Number of leg-biased genes in males under poor (P) or rich (R) nutrition. L1- and L3-biased genes are represented in yellow and blue respectively.

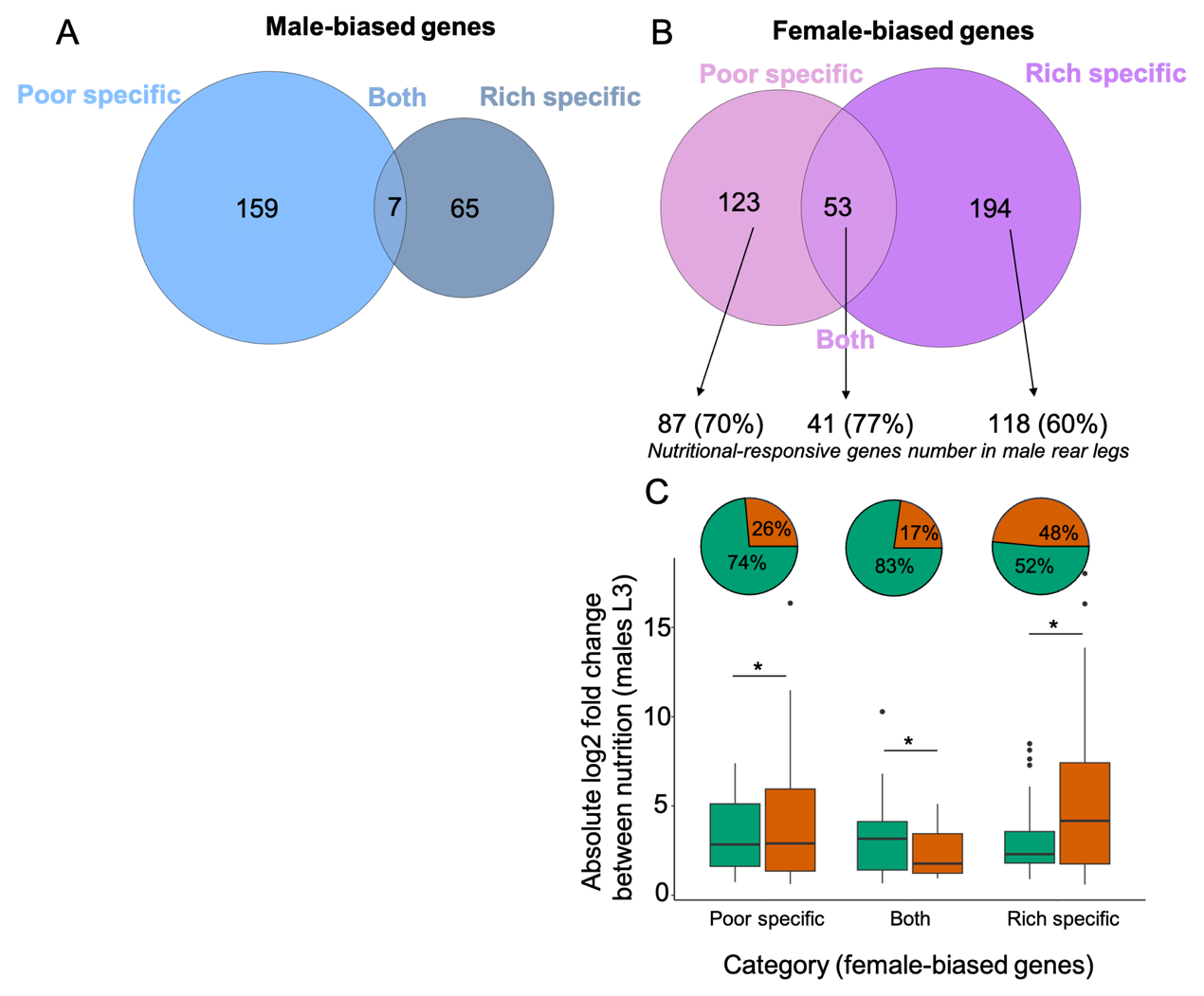

**Supplementary figure S7**: (**A, B)** Sex-biased genes according to nutritional condition (**A)** Male-biased genes identified under poor and/or rich nutrition. (**B**) Female-biased genes identified specifically under poor nutrition (poor specific), under both nutritional condition (both), or specifically under rich nutrition (rich specific). For each category, the number and percentage of genes that were also nutrition-responsive in male rear legs are indicated. (**C**) Direction and magnitude of the nutritional response of these female-biased, nutrition-responsive genes in male rear legs. Pie charts show the proportion of poor- and rich-biased genes (green and orange, respectively) within each category of female bias. Bowplot show the absolute log2 fold change between nutritional condition. Differences between poor- and rich-biased genes within each category were assessed using Wilcoxon rank-sum tests (* : p < 0.05).

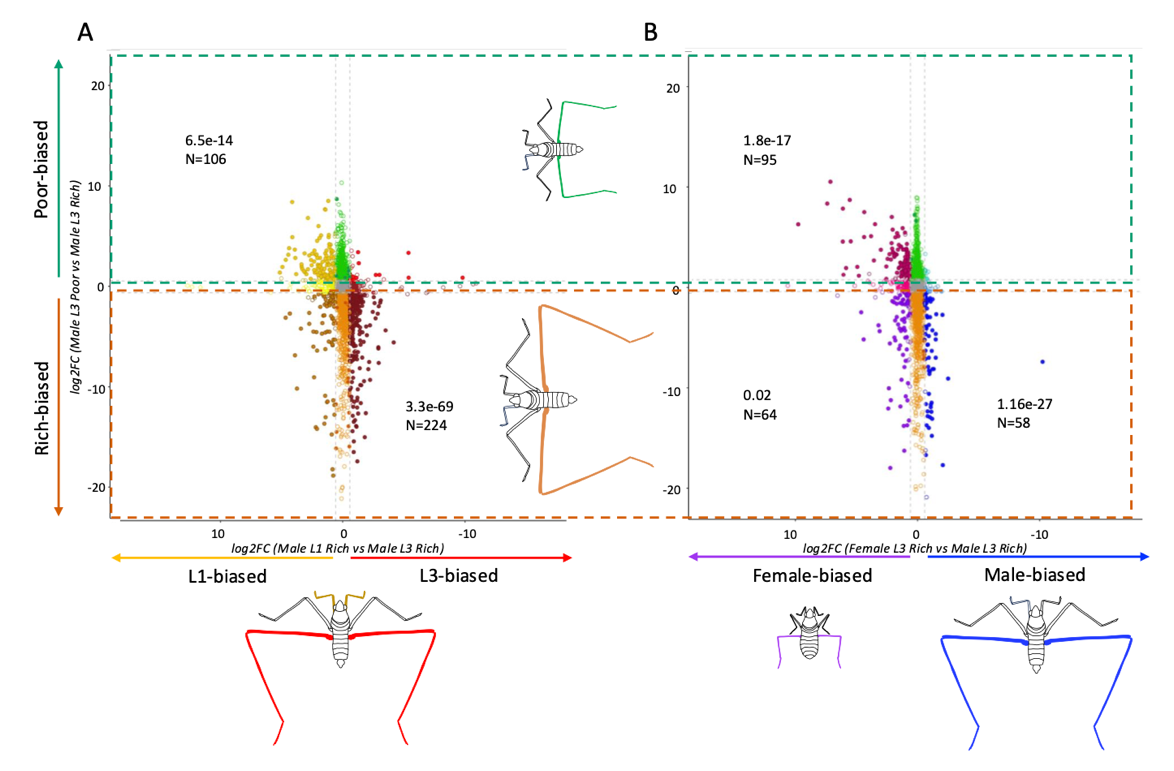

**Supplementary figure S8**: (**A, B)** Correlation profiles of nutrition-biased genes in male rear legs (y axis) and (e) leg-biased genes in rich male or (f) sex-biased genes in rich individuals. Unbiased genes are either grey (no biased), orange or green are rich- or poor-biased genes in L3M that are not found in leg- or sex-biased genes in rich individuals. Filled circles indicate genes with padj < 0.05 in both conditions. Hollow circles indicate genes with padj > 0.05 in one or both conditions.  **e** Yellow or red filled circles are poor-biased genes in L3M that are also either L1- or L3-biased respectively. Dark yellow or brown filler circles are rich-biased genes in L3M that are either L1- or L3-biased respectively. **f** Pink or turquoise filled genes are poor-biased genes that are either female- or male-biased respectively. Purple or dark blue filled circles are rich-biased genes in L3M that are either female- or male-biased respectively. Enrichment of leg- or sex-biased genes in poor and rich males was estimated using the Fisher’s exact tests, where significative enrichments (p-value<0.05) are referred in the panel.

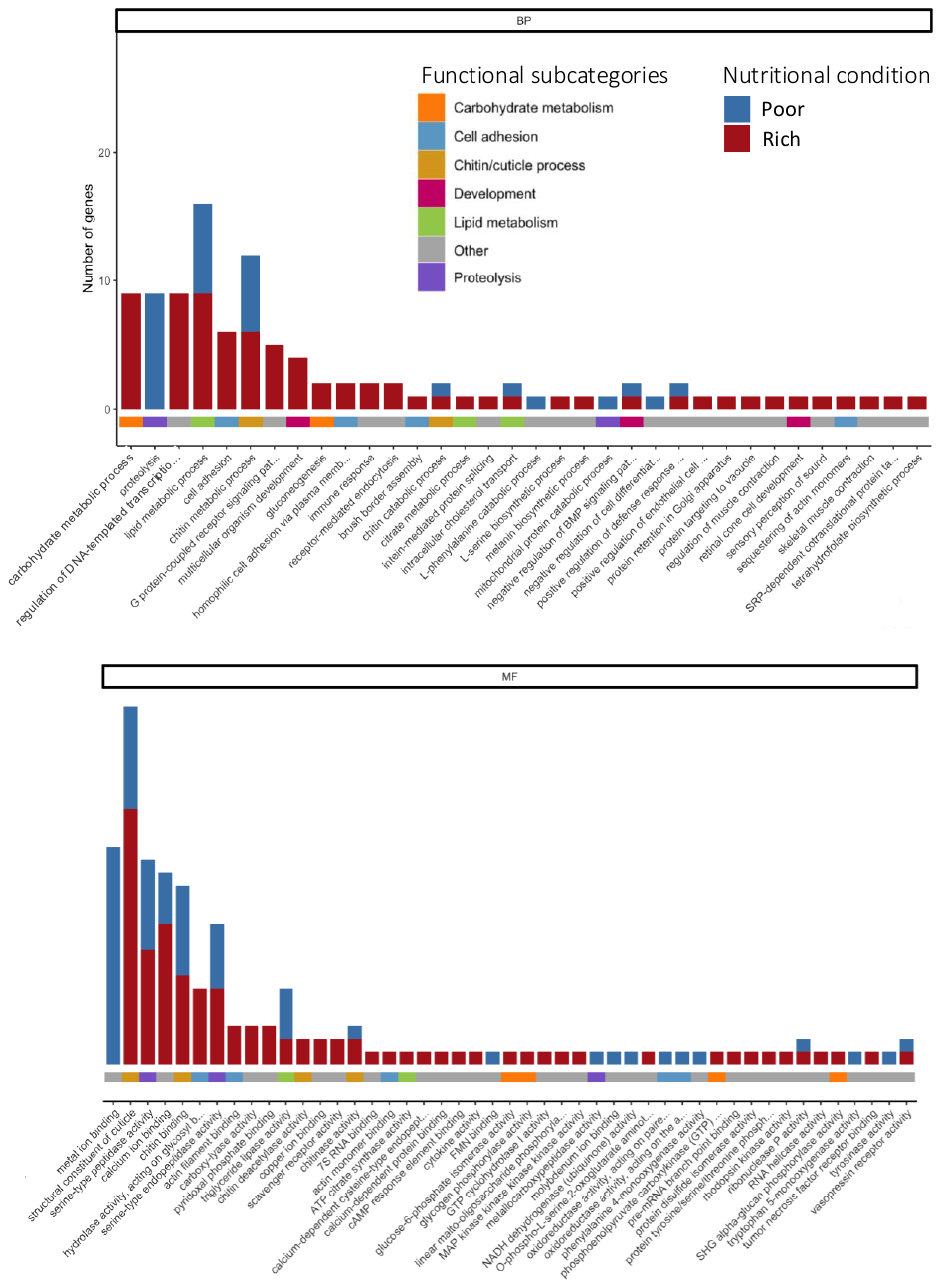

**Supplementary figure S9**: Gene ontology (GO) Analysis in L3-biased genes under poor or rich nutrition (blue and red bar respectively). Significant gene number from significant Term (classicFisher <0.05) are represented in y axis. GO terms from Biological Process or Molecular Function (BP and MF respectively) categories are represented in the x axis. GO terms are grouped into functional subcategories, which are represented by colors under histograms.

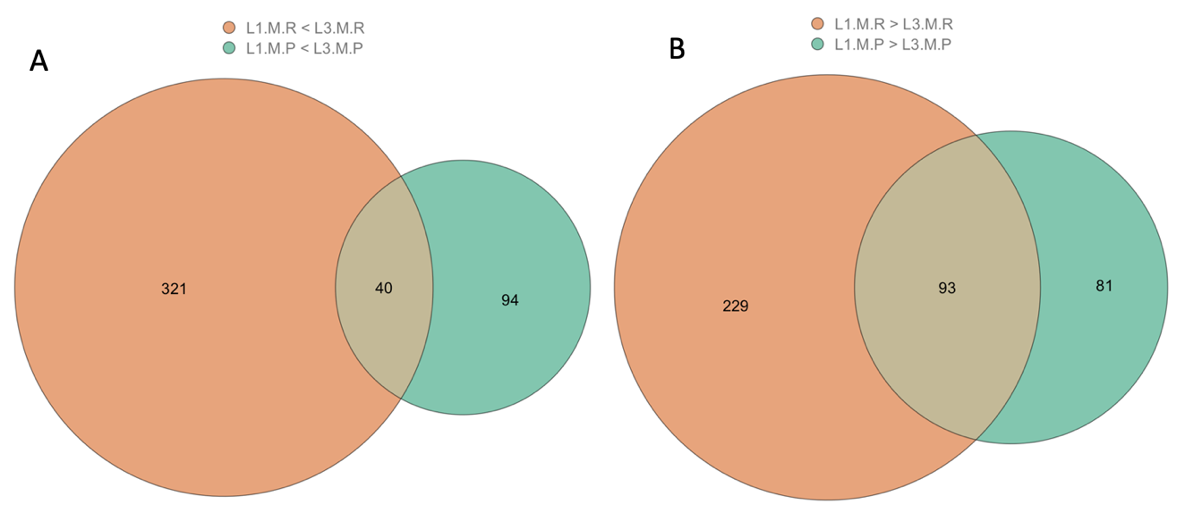

**Supplementary figure S10**: (**A, B)** Leg-biased genes according to nutritional condition: (a**)** L1-biased and (b) L3-biased genes identified in males under poor and/or rich nutrition (green, mixed color and orange, respectively).

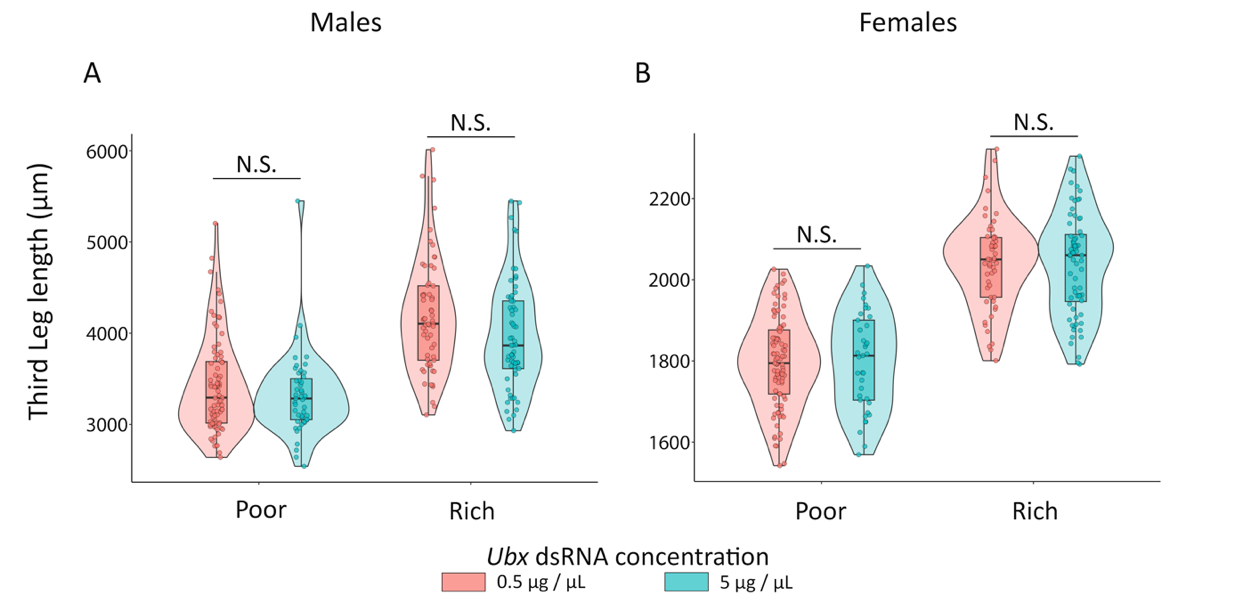

**Supplementary figure S11**: Knock-down of *Ubx* in *Microvelia longipes* long-legged isogenic line. *Ubx* dsRNA was injected in third instar nymphs using two concentrations, 0.5 µg/µL and 5 µg/µL (pink and blue, respectively). Nymphs were treated by poor or rich nutrition after injection. We measured rear leg length in injected (a) males and (b) females at adult stage. We did not see differences between the two concentrations (two-way ANOVA, p-value >0.05).
